## Supplementary figures and images for "Optogenetic silencing hippocampal inputs to the retrosplenial cortex causes a prolonged disruption of spatial working memory"

### Supplementary Figure S1

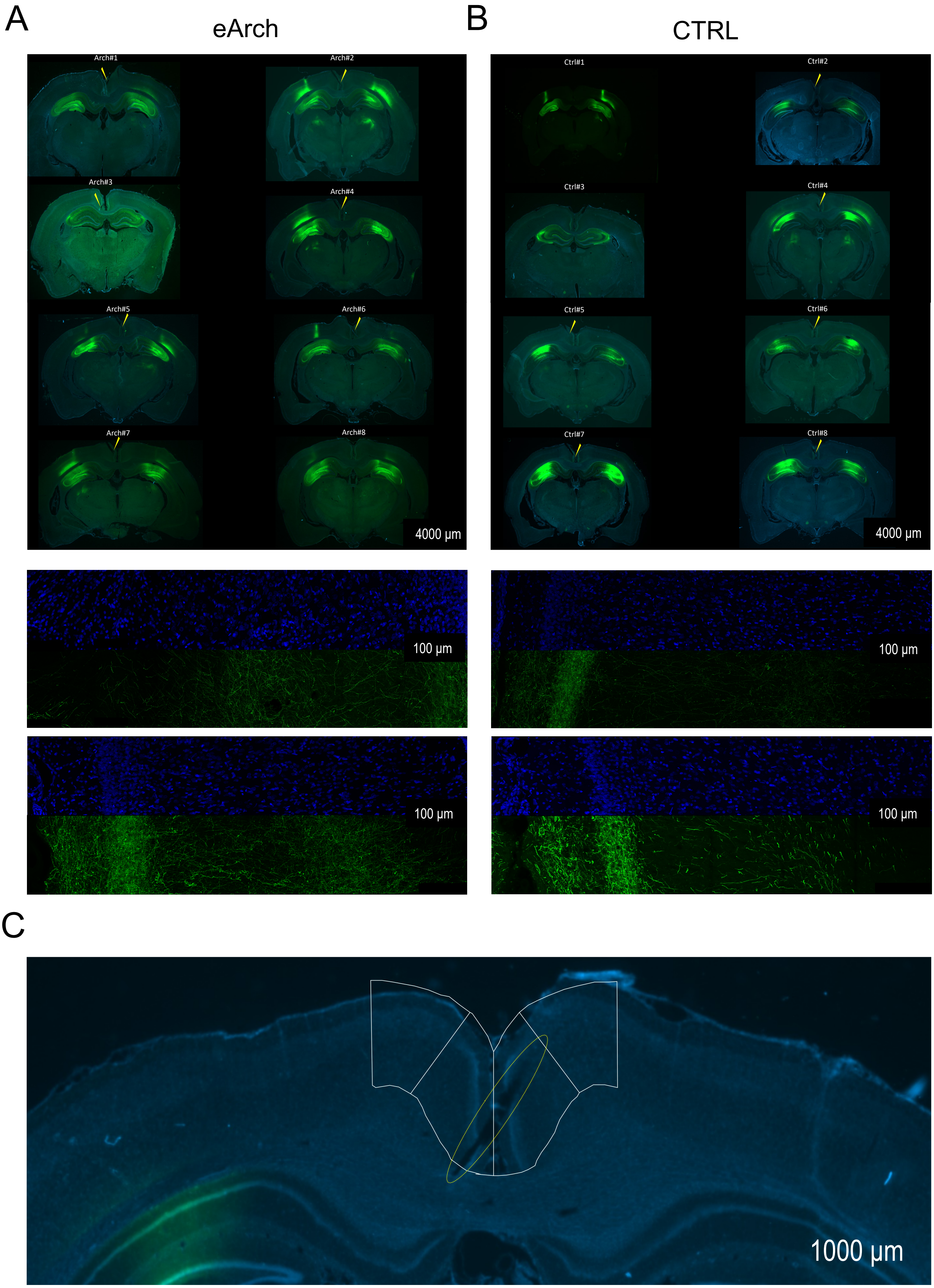
