## Supplementary GLMM Tables 1-8 for "Optogenetic silencing hippocampal inputs to the retrosplenial cortex causes a prolonged disruption of spatial working memory"

**Table 1. Generalized Linear Mixed Model analysis comparing performance between eArch+ and CTRL, observed in baseline sessions (corresponding to the plots in figure 2C)**

| Model Information | |
|---|---|
| **Function** | outcome ~ 1 + group + (1\|rat) |
| **Link function** | Logit |
| **Distribution** | Binomial |

Fixed Effects Parameter Estimates

| | | | | | 95% Exp(B) Confidence Interval | | | |
|---|---|---|---|---|---|---|---|---|
| **Names** | **Effect** | **Estimate** | **SE** | **exp(B)** | **Lower** | **Upper** | **z** | **p** |
| (Intercept) | (Intercept) | 1.653 | 0.0761 | 5.22 | 4.500 | 6.06 | 21.71 | < .001 |
| group1 | CTRL - ARCH | 0.190 | 0.1523 | 1.21 | 0.897 | 1.63 | 1.25 | 0.212 |

Fixed Effect Omnibus tests

| | $X^2$ | df | p |
|---|---|---|---|
| group | 1.56 | 1.00 | 0.212 |

Post Hoc Comparisons - group

| Comparison | | | | | | |
|---|---|---|---|---|---|---|
| **group** | | **group** | **exp(B)** | **SE** | **z** | $p_{bonferroni}$ |
| ARCH | - | CTRL | 0.827 | 0.126 | -1.25 | 0.212 |

### Table 2. Generalized Linear Mixed Model analysis comparing the ratio of correct trials between eArch+ and CTRL groups, and illuminated and non-illuminated trials (corresponding to the plots in figure 3A and 3B)

Model Information

| | |
|---|---|
| **Function** | outcome ~ 1 + illumination + group + illumination:group + (1\|rat) |
| **Link function** | Logit |
| **Distribution** | Binomial |

Fixed Effects Parameter Estimates

| | | | | | 95% Exp(B) Confidence Interval | | | |
|---|---|---|---|---|---|---|---|---|
| **Names** | **Effect** | **Estimate** | **SE** | **exp(B)** | **Lower** | **Upper** | **z** | **p** |
| (Intercept) | (Intercept) | 1.2265 | 0.0762 | 3.409 | 2.936 | 3.96 | 16.087 | < .001 |
| illumination1 | 1 - 0 | 0.0120 | 0.0721 | 1.012 | 0.879 | 1.17 | 0.166 | 0.868 |
| group1 | CTRL - ARCH | 0.4198 | 0.1524 | 1.522 | 1.129 | 2.05 | 2.755 | 0.006 |
| illumination1 ✻ group1 | 1 - 0 ✻ CTRL - ARCH | -0.0547 | 0.1442 | 0.947 | 0.714 | 1.26 | -0.379 | 0.704 |

Fixed Effect Omnibus tests

| | $X^2$ | df | p |
|---|---|---|---|
| illumination | 0.0275 | 1.00 | 0.868 |
| group | 7.5923 | 1.00 | 0.006 |
| illumination ✻ group | 0.1439 | 1.00 | 0.704 |

Post Hoc Comparisons - group

| Comparison | | | | | | |
|---|---|---|---|---|---|---|
| **group** | | **group** | **exp(B)** | **SE** | **z** | **$p_{bonferroni}$** |
| ARCH | - | CTRL | 0.657 | 0.100 | -2.76 | 0.006 |

Post Hoc Comparisons - illumination

| Comparison | | | | | | |
|---|---|---|---|---|---|---|
| **illumination** | | **illumination** | **exp(B)** | **SE** | **z** | **$p_{bonferroni}$** |
| 0 | - | 1 | 0.988 | 0.0712 | -0.166 | 0.868 |

**Commented [BP1]:** This p-value corresponds to figure 3A

**Commented [BP2]:** In my revision of the manuscript I commented that the legend of figure 3B was unclear. I'm not sure what you ended up writing as the legend but in tables 2 and 3 I highlighted the p-values that could've been included

Simple effects of group : Parameter estimates

| Moderator levels | | | | | 95% Exp(B) Confidence Interval | | | |
|---|---|---|---|---|---|---|---|---|
| illumination | contrast | Estimate | SE | exp(B) | Lower | Upper | z | p |
| 0 | CTRL - ARCH | 0.447 | 0.168 | 1.56 | 1.12 | 2.18 | 2.66 | 0.008 |
| 1 | CTRL - ARCH | 0.392 | 0.169 | 1.48 | 1.06 | 2.06 | 2.33 | 0.020 |

Simple effects of group : Omnibus Tests

| Moderator levels | | | |
|---|---|---|---|
| illumination | X² | df | p |
| 0 | 7.06 | 1.00 | 0.008 |
| 1 | 5.41 | 1.00 | 0.020 |

### Table 3. Generalized Linear Mixed Model analysis comparing the ratio of correct trials between baseline, illuminated and non-illuminated trials, for eArch+ and CTRL groups (corresponding to the plots in figure 3B)

Model Information

| | |
|---|---|
| **Function** | outcome ~ 1 + illumination_epoch + group + illumination_epoch:group + (1\|rat) |
| **Link function** | Logit |
| **Distribution** | Binomial |

Legend of illumination epochs: 0 – non-illuminated, 1 – illuminated trial, 2 - baseline

Fixed Effects Parameter Estimates

| | | | | | 95% Exp(B) Confidence Interval | | | |
|---|---|---|---|---|---|---|---|---|
| **Names** | **Effect** | **Estimate** | **SE** | **exp(B)** | **Lower** | **Upper** | **z** | **p** |
| (Intercept) | (Intercept) | 1.3721 | 0.0683 | 3.944 | 3.450 | 4.51 | 20.090 | < .001 |
| group1 | CTRL - ARCH | 0.3431 | 0.1364 | 1.409 | 1.079 | 1.84 | 2.515 | 0.012 |
| illum_epoch1 | 1 - 0 | 0.0120 | 0.0720 | 1.012 | 0.879 | 1.17 | 0.166 | 0.868 |
| illum_epoch2 | 2 - 0 | 0.4562 | 0.0917 | 1.578 | 1.318 | 1.89 | 4.975 | < .001 |
| group1 ✻ illum_epoch1 | CTRL - ARCH ✻ 1 - 0 | -0.0541 | 0.1440 | 0.947 | 0.714 | 1.26 | -0.376 | 0.707 |
| group1 ✻ illum_epoch2 | CTRL - ARCH ✻ 2 - 0 | -0.2785 | 0.1833 | 0.757 | 0.528 | 1.08 | -1.519 | 0.129 |

Fixed Effect Omnibus tests

| | $X^2$ | df | p |
|---|---|---|---|
| group | 6.32 | 1.00 | 0.012 |
| Illum_epoch | 28.45 | 2.00 | < .001 |
| group ✻ illum_epoch | 2.36 | 2.00 | 0.307 |

Post Hoc Comparisons - group ✻ illumination epoch

| Comparison | | | | | | | | |
|---|---|---|---|---|---|---|---|---|
| **group** | **Illum_epoch** | | **group** | **Illum_epoch** | **exp(B)** | **SE** | **z** | $p_{bonferroni}$ |
| ARCH | 0 | - | ARCH | 1 | 0.962 | 0.0942 | -0.398 | 1.000 |
| ARCH | 0 | - | ARCH | 2 | 0.551 | 0.0698 | -4.701 | < .001 |
| ARCH | 0 | - | CTRL | 0 | 0.635 | 0.0984 | -2.930 | 0.051 |
| ARCH | 0 | - | CTRL | 1 | 0.645 | 0.0999 | -2.833 | 0.069 |
| ARCH | 0 | - | CTRL | 2 | 0.463 | 0.0807 | -4.419 | < .001 |
| ARCH | 1 | - | ARCH | 2 | 0.573 | 0.0730 | -4.370 | < .001 |
| ARCH | 1 | - | CTRL | 1 | 0.670 | 0.1042 | -2.573 | 0.151 |

Post Hoc Comparisons - group ✻ illumination epoch

| Comparison | | | | | | | | |
|---|---|---|---|---|---|---|---|---|
| group | Illum_epoch | | group | Illum_epoch | exp(B) | SE | z | $p_{bonferroni}$ |
| ARCH | 1 | - | CTRL | 2 | 0.481 | 0.0841 | -4.185 | < .001 |
| ARCH | 2 | - | CTRL | 2 | 0.839 | 0.1615 | -0.912 | 1.000 |
| CTRL | 0 | - | ARCH | 1 | 1.514 | 0.2355 | 2.669 | 0.114 |
| CTRL | 0 | - | ARCH | 2 | 0.868 | 0.1519 | -0.808 | 1.000 |
| CTRL | 0 | - | CTRL | 1 | 1.015 | 0.1071 | 0.143 | 1.000 |
| CTRL | 0 | - | CTRL | 2 | 0.728 | 0.0966 | -2.391 | 0.252 |
| CTRL | 1 | - | ARCH | 2 | 0.855 | 0.1496 | -0.894 | 1.000 |
| CTRL | 1 | - | CTRL | 2 | 0.717 | 0.0951 | -2.505 | 0.184 |

**Table 4. Generalized Linear Mixed Model analysis comparing the probability of the current trial being correct following an illuminated trial (corresponding to the plots in figure 3C)**

Model Information

| | |
|---|---|
| **Function** | Future_outcome ~ 1 + group + outcome + group:outcome + (1\|rat) |
| **Link function** | Logit |
| **Distribution** | Binomial |

Fixed Effects Parameter Estimates

| | | | | | 95% Exp(B) Confidence Interval | | | |
|---|---|---|---|---|---|---|---|---|
| **Names** | **Effect** | **Estimate** | **SE** | **exp(B)** | **Lower** | **Upper** | **z** | **p** |
| (Intercept) | (Intercept) | 1.1117 | 0.0862 | 3.039 | 2.567 | 3.60 | 12.9022 | < .001 |
| group1 | CTRL - ARCH | 0.4175 | 0.1717 | 1.518 | 1.084 | 2.13 | 2.4308 | 0.015 |
| outcome1 | 1 - 0 | 0.0539 | 0.1219 | 1.055 | 0.831 | 1.34 | 0.4425 | 0.658 |
| group1 ✻ outcome1 | CTRL - ARCH ✻ 1 - 0 | -0.0215 | 0.2431 | 0.979 | 0.608 | 1.58 | -0.0885 | 0.929 |

Fixed Effect Omnibus tests

| | $X^2$ | df | p |
|---|---|---|---|
| group | 5.90903 | 1.00 | 0.015 |
| outcome | 0.19579 | 1.00 | 0.658 |
| group ✻ outcome | 0.00783 | 1.00 | 0.929 |

Post Hoc Comparisons - group

| Comparison | | | | | | |
|---|---|---|---|---|---|---|
| **group** | | **group** | **exp(B)** | **SE** | **z** | $p_{bonferroni}$ |
| ARCH | - | CTRL | 0.659 | 0.113 | -2.43 | 0.015 |

### Table 5. Generalized Linear Mixed Model analysis comparing the probability of errors in the current trial following an error trial (corresponding to the plots in figure 4A)

Model Information

| | |
|---|---|
| **Function** | Errors ~1 + group + illumination + group:illumination + (1\|session_number) + (1\|run_number) |
| **Link function** | Logit |
| **Distribution** | Binomial |

Fixed Effects Parameter Estimates

| | | | | | 95% Exp(B) Confidence Interval | | | |
|---|---|---|---|---|---|---|---|---|
| **Names** | **Effect** | **Estimate** | **SE** | **exp(B)** | **Lower** | **Upper** | **z** | **p** |
| (Intercept) | (Intercept) | -1.239 | 0.107 | 0.290 | 0.235 | 0.357 | -11.60 | < .001 |
| group1 | CTRL - ARCH | -0.593 | 0.159 | 0.553 | 0.404 | 0.755 | -3.72 | < .001 |
| Illumination1 | 1 - 0 | 0.160 | 0.156 | 1.173 | 0.865 | 1.592 | 1.03 | 0.305 |
| group1 ✻ illumination1 | CTRL - ARCH ✻ 1 - 0 | 0.325 | 0.311 | 1.384 | 0.752 | 2.545 | 1.04 | 0.296 |

Fixed Effect Omnibus tests

| | $X^2$ | df | p |
|---|---|---|---|
| group | 13.84 | 1.00 | < .001 |
| illumination | 1.05 | 1.00 | 0.305 |
| group ✻ illumination | 1.09 | 1.00 | 0.296 |

Post Hoc Comparisons - group

| Comparison | | | | | | |
|---|---|---|---|---|---|---|
| **group** | | **group** | **exp(B)** | **SE** | **z** | $p_{bonferroni}$ |
| ARCH | - | CTRL | 1.81 | 0.289 | 3.72 | < .001 |

**Table 6. Generalized Linear Mixed Model analysis comparing the probability of errors in the current trial following an error trial, in baseline sessions (corresponding to the plots in figure 4B)**

Model Information

| | |
|---|---|
| **Function** | Errors ~1 + group + (1\|session_number) + (1\|run_number) |
| **Link function** | Logit |
| **Distribution** | Binomial |

Fixed Effects Parameter Estimates

| | | | | | 95% Exp(B) Confidence Interval | | | |
|---|---|---|---|---|---|---|---|---|
| **Names** | **Effect** | **Estimate** | **SE** | **exp(B)** | **Lower** | **Upper** | **z** | **p** |
| (Intercept) | (Intercept) | -2.133 | 0.375 | 0.118 | 0.0567 | 0.247 | -5.68 | < .001 |
| group1 | CTRL - ARCH | -0.622 | 0.456 | 0.537 | 0.2197 | 1.311 | -1.37 | 0.172 |

Fixed Effect Omnibus tests

| | **X²** | **df** | **p** |
|---|---|---|---|
| group | 1.86 | 1.00 | 0.172 |

Post Hoc Comparisons - group

| **Comparison** | | | | | | |
|---|---|---|---|---|---|---|
| **group** | | **group** | **exp(B)** | **SE** | **z** | $p_{bonferroni}$ |
| ARCH | - | CTRL | 1.86 | 0.849 | 1.37 | 0.172 |

### Table 7. Generalized Linear Mixed Model analysis comparing the probability of correct trials following illumination of trial$_{T-n}$ (corresponding to the plots in figure 5A)

Model Information

| | |
|---|---|
| **Function** | $trial_{n-x}$ ~1 + group + (1\|rat) |
| **Link function** | Logit |
| **Distribution** | Binomial |

Fixed Effects Parameter Estimates

| | | | | | | 95% Exp(B) Confidence Interval | | | |
|---|---|---|---|---|---|---|---|---|---|
| | **Names** | **Effect** | **Estimate** | **SE** | **exp(B)** | **Lower** | **Upper** | **z** | **p** |
| t-1 | (Intercept) | (Intercept) | 1.137 | 0.0795 | 3.12 | 2.67 | 3.64 | 14.30 | < .001 |
| | group1 | ctrl - arch | 0.406 | 0.1586 | 1.50 | 1.10 | 2.05 | 2.56 | 0.010 |
| t-2 | (Intercept) | (Intercept) | 1.161 | 0.0949 | 3.19 | 2.65 | 3.84 | 12.23 | < .001 |
| | group1 | ctrl - arch | 0.403 | 0.1894 | 1.50 | 1.03 | 2.17 | 2.13 | 0.033 |
| t-3 | (Intercept) | (Intercept) | 1.288 | 0.0799 | 3.63 | 3.10 | 4.24 | 16.12 | < .001 |
| | group1 | ctrl - arch | 0.334 | 0.1590 | 1.40 | 1.02 | 1.91 | 2.10 | 0.036 |
| t-4 | (Intercept) | (Intercept) | 1.122 | 0.0939 | 3.07 | 2.554 | 3.69 | 11.95 | < .001 |
| | group1 | ctrl - arch | 0.330 | 0.1872 | 1.39 | 0.964 | 2.01 | 1.76 | 0.078 |

Fixed Effect Omnibus tests

| | | $X^2$ | df | p |
|---|---|---|---|---|
| t-1 | group | 6.56 | 1.00 | 0.010 |
| t-2 | group | 4.52 | 1.00 | 0.033 |
| t-3 | group | 4.42 | 1.00 | 0.036 |
| t-4 | group | 3.11 | 1.00 | 0.078 |

Post Hoc Comparisons - group

| | Comparison | | | | | | |
|---|---|---|---|---|---|---|---|
| | **group** | | **group** | **exp(B)** | **SE** | **z** | $p_{bonferroni}$ |
| t-1 | arch | - | ctrl | 0.666 | 0.106 | -2.56 | 0.010 |

Post Hoc Comparisons - group

| Comparison | | | | | | | |
|---|---|---|---|---|---|---|---|
| | group | | group | exp(B) | SE | z | $p_{bonferroni}$ |
| t-2 | arch | - | ctrl | 0.669 | 0.127 | -2.13 | 0.033 |
| t-3 | arch | - | ctrl | 0.716 | 0.114 | -2.10 | 0.036 |
| t-4 | arch | - | ctrl | 0.719 | 0.135 | -1.76 | 0.078 |

### Table 8. Generalized Linear Mixed Model analysis comparing the log(time) spent at the choice point (corresponding to the plots in figure 6A)

| Model Information | |
|---|---|
| **Function** | Normalized_log(time) ~1 + illumination + group + illumination:group + (1\|rat) |
| **Link function** | Identity |
| **Distribution** | Gaussian |

Fixed Effects Parameter Estimates

| | | | | 95% Confidence Interval | | | | |
|---|---|---|---|---|---|---|---|---|
| **Names** | **Effect** | **Estimate** | **SE** | **Lower** | **Upper** | **df** | **t** | **p** |
| (Intercept) | (Intercept) | -0.1971 | 0.248 | -0.683 | 0.289 | 13.8 | -0.795 | 0.440 |
| group1 | CTRL - ARCH | 1.2674 | 0.496 | 0.295 | 2.239 | 13.8 | 2.556 | 0.023 |
| illumination1 | 1 - 0 | 0.0468 | 0.274 | -0.491 | 0.585 | 4249.3 | 0.171 | 0.865 |
| group1 ✻ illumination 1 | CTRL - ARCH ✻ 1 - 0 | -0.6153 | 0.549 | -1.691 | 0.460 | 4249.3 | -1.121 | 0.262 |

Fixed Effect Omnibus tests

| | F | Num df | Den df | p |
|---|---|---|---|---|
| group | 6.5315 | 1 | 13.8 | 0.023 |
| illumination | 0.0291 | 1 | 4249.3 | 0.865 |
| group ✻ illumination | 1.2571 | 1 | 4249.3 | 0.262 |

Nota. Satterthwaite method for degrees of freedom

Table 9. Generalized Linear Mixed Model of the log(time) spent at the choice point by illumination and outcome (corresponding to plot of figure 6B)

| Model Information | |
|---|---|
| **Function** | Normalized_log(time) ~1 + illumination + group + outcome + illumination:group + illumination:outcome + group:outcome + illumination:group:outcome + (1\|rat) |
| **Link function** | Identity |
| **Distribution** | Gaussian |

Fixed Effects Parameter Estimates

| Names | Effect | Estimate | SE | 95% Confidence Interval Lower | 95% Confidence Interval Upper | df | t | p |
|---|---|---|---|---|---|---|---|---|
| (Intercept) | (Intercept) | -0.3571 | 0.264 | -0.874 | 0.159 | 17.9 | -1.3550 | 0.192 |
| illumination 1 | 1 - 0 | 0.0277 | 0.330 | -0.619 | 0.675 | 4246.3 | 0.0840 | 0.933 |
| group1 | CTRL - ARCH | 0.9148 | 0.527 | -0.118 | 1.948 | 17.9 | 1.7355 | 0.100 |
| outcome1 | 1 - 0 | 0.5136 | 0.332 | -0.137 | 1.164 | 4255.0 | 1.5471 | 0.122 |
| illumination 1 ✻ group1 | 1 - 0 ✻ CTRL - ARCH | -0.8037 | 0.660 | -2.098 | 0.491 | 4246.3 | -1.2171 | 0.224 |
| illumination 1 ✻ outcome1 | 1 - 0 ✻ 1 - 0 | 0.0323 | 0.661 | -1.262 | 1.327 | 4248.6 | 0.0489 | 0.961 |
| group1 ✻ outcome1 | CTRL - ARCH ✻ 1 - 0 | 1.1575 | 0.664 | -0.144 | 2.459 | 4255.0 | 1.7432 | 0.081 |
| illumination1 ✻ group1 ✻ outcome1 | 1 - 0 ✻ CTRL - ARCH ✻ 1 - 0 | 0.6806 | 1.321 | -1.909 | 3.270 | 4248.6 | 0.5152 | 0.606 |

Fixed Effect Omnibus tests

| | F | Num df | Den df | p |
|---|---|---|---|---|
| illumination | 0.00705 | 1 | 4246.3 | 0.933 |
| group | 3.01211 | 1 | 17.9 | 0.100 |
| outcome | 2.39345 | 1 | 4255.0 | 0.122 |
| illumination ✻ group | 1.48133 | 1 | 4246.3 | 0.224 |
| illumination ✻ outcome | 0.00239 | 1 | 4248.6 | 0.961 |
| group ✻ outcome | 3.03866 | 1 | 4255.0 | 0.081 |
| illumination ✻ group ✻ outcome | 0.26539 | 1 | 4248.6 | 0.606 |

Nota. Satterthwaite method for degrees of freedom

Simple effects of group : Parameter estimates

| Moderator levels | | | | | 95% Confidence Interval | | | | |
|---|---|---|---|---|---|---|---|---|---|
| illum | outcome | contrast | Estimate | SE | Lower | Upper | df | t | p |
| 0 | 0 | CTRL - ARCH | 0.908 | 0.915 | -0.8992 | 2.72 | 157.8 | 0.992 | 0.323 |
| | 1 | CTRL - ARCH | 1.725 | 0.603 | 0.4949 | 2.96 | 30.5 | 2.862 | 0.008 |
| 1 | 0 | CTRL - ARCH | -0.236 | 0.927 | -2.0654 | 1.59 | 166.3 | -0.255 | 0.799 |
| | 1 | CTRL - ARCH | 1.262 | 0.604 | 0.0290 | 2.49 | 30.9 | 2.088 | 0.045 |

Simple effects of group : Omnibus Tests

| Moderator levels | | | | | |
|---|---|---|---|---|---|
| illumination | outcome | F | Num df | Den df | p |
| 0 | 0 | 0.9850 | 1.00 | 157.8 | 0.323 |
| | 1 | 8.1900 | 1.00 | 30.5 | 0.008 |
| 1 | 0 | 0.0650 | 1.00 | 166.3 | 0.799 |
| | 1 | 4.3580 | 1.00 | 30.9 | 0.045 |
